# Evidence of flowering time advance in blue lupin (*Lupinus angustifolius*) in the last decades revealed by herbarium data and citizen science databases

**DOI:** 10.1101/2024.12.21.628886

**Authors:** F.J. Jiménez – López, C. Lara – Romero, J.M. Iriondo, M.L. Rubio Teso, A García – Fernández

## Abstract

Rapid adaptation to climate change in plants manifests through genetic modifications, epigenetic changes, or species-microbiome interactions, with short-living species having a greater potential for adaptation. The integration of historical collection data with modern databases improves our ability to study plant phenology and distribution shifts in response to global change. This study evaluates changes in the flowering phenology and vernalization requirements of *Lupinus angustifolius* populations across the Iberian Peninsula, assessing the effectiveness of diverse data sources. Using herbarium specimens and field photographs, we compiled flowering data and estimated flowering peaks. Thermal time and vernalization requirements were estimated using recorded flowering stages and high-resolution climate data. Analyses revealed a progressive advancement in *L. angustifolius* flowering in the last 60 years. Latitude and elevation emerged as significant influencing factors, with greater shifts observed at lower latitudes and elevations. Notably, plants now flower with fewer vernalization days, and higher thermal time, particularly at lower elevations. The adaptive responses of plants to global change are intricate and dynamic. Phenological advancement in flowering appears to be a key strategy for coping with environmental alterations, facilitating survival and reproduction through the interplay of genotype, phenotype, and environment, as well as existing genetic diversity. In this context, historical records and long-term monitoring prove invaluable for assessing the efficacy and pace of these adaptive processes. To improve our understanding of plant adaptation in the context of climate change, it is essential to synthesize diverse data sources and maintain ongoing collection efforts.

## INTRODUCCTION

Rapid adaptation has been proposed as an essential mechanism for several species when they try to achieve adaptative responses against climate change (Shaw & Etterson, 2012). Fast adaptation can be led by genome modifications or allele frequency fluctuations, but can also be modulated by epigenetic change or species-microbiome interactions (Torda et al., 2017). The trade-offs between these factors influence the rate at which evolution occurs, and can vary across species (Bromham, 2009). Short-living species, such as annual plants or insects, that can produce one (or more) generations per season, are more prone to generate new genotypes that foster adaptation processes (Kristensen et al., 2018). On the other hand, long-living species have more difficulties to prompt these changes, due to generation overlapping or genotype mixture (Friedman, 2020; but also see Whittle & Johnson, 2003). Other variables, such as unequal environmental pressures or specific trade-offs between traits can also influence the rates of adaptation between populations or species (Kristensen et al., 2018).

Flowering phenology has been recently described as one of the most disturbed traits in the current context of climate change (Inouye, 2022; Valdés et al., 2023; Collins et al., 2024). Shifts in flowering time may modify plant-pollinator interactions (Morente-López et al. 2018), with important consequences for population fitness and fecundity (Petanidou et al., 2014; Duchenne et al., 2020). Advancing flowering onset or modifying the flowering duration may alter local population adaptation and thus, evolutionary resilience in response to global change (Anderson et al., 2012; Ramirez-Parada et al., 2024). Furthermore, flowering phenology fluctuations could also impact plant restoration efforts, as these shifts may disrupt critical ecological interactions and processes necessary for successful restoration. Temporal series and long-term monitoring of flowering periods are crucial to understanding the mechanisms driving these shifts, and such knowledge can inform strategies for restoration and conservation by predicting and mitigating phenological mismatches (Gordo & Sanz, 2009; Langvall & Ottosson Löfvenius, 2021). The processes that govern flowering phenology are complex and tangled, including gene expression, physiological pathways and plant-plant or plant-microbiome interactions (Elzinga et al., 2007; Kudoh, 2016). Among these processes, vernalization plays a pivotal role in many species, acting as a mechanism by which plants need to be exposed to winter cold temperatures to develop subsequent flowering structures (Chouard, 1960). This mechanism is a common patway in several plant families, suggesting a convergent evolutionary process and the implication of different genes and metabolic routes during its regulation (Ream et al., 2012). On the other hand, thermal time, or the accumulation of heat units above a base temperature, is crucial for the timing of flowering (Li et al. 2014; Capovilla et al. 2015). Plants require a specific amount of thermal time to transition from vegetative growth to flowering, which varies among species and is influenced by environmental conditions. The relevance of these mechanism also involves population and community dynamics, modulating flowering synchrony between individuals (Meyer et al., 2004; Huang et al., 2024) or plant-pollinator interactions (Forrest & Thomson, 2011). The expected shift in the temperatures, associated to climate change, may modify vernalization thresholds for plant species. Changes in these thresholds suggest that plants will need to develop new adaptation processes to cope with the changing climate.

Different plant responses to global change (e.g. phenotypic plasticity, local adaptation) might trigger shifts in plant phenology (Anderson et al., 2012; Inouye, 2022), including all processes related with flowering. Under semiarid and Mediterranean environments with severe drought conditions, flowering onset has advanced to encompass flowering peak and duration with water availability and avoid summer abiotic stress (Pareja-Bonilla et al., 2023). However, the response of plants to these conditions can vary significantly depending on the species and specific environmental factors, making it a complex process to predict and generalize (Crimmins et al., 2010; Kazan & Lyons, 2016). On the other hand, in humid ecosystems, such as mountain or alpine habitats, plant responses related with flowering show different fluctuations to encompass the timing of snow melt, extreme events and summer droughts, which affect plant fitness (Rafferty et al., 2020; Sethi et al., 2020; Zu et al., 2023).

Traditionally, herbarium material has been employed for taxonomic descriptions, biogeographic and phylogeography studies and, if possible, as source material for genetic approaches (e.g. Lister et al., 2010; Marsico et al., 2020). Nowadays, herbarium sheets are widely considered for other studies, and have been used, for example, to evaluate the changes in plant traits during an extended period, or the accumulation of toxics in plant tissues, (Belloeil et al., 2021; Purwadi et al., 2021). Furthermore, herbarium specimens are increasingly used to evaluate temporal changes in flowering, spatial distributions, or colonization-extinction events (Jones & Daehler, 2018; Bates et al., 2023; Rondinel-Mendoza et al., 2024). Therefore, collections data provide an interesting framework of study that must be integrated in ecological-evolutionary-biological studies.

Integrating collections data with current global biodiversity databases (including social networks and citizen science platforms) can provide complementary information at different spatial and temporal scales. Citizen science databases represent a modern and non-exclusive contribution to the records in scientific collection datasets. They are available to anyone, allowing a significant increase in the information collected on the distribution or phenology of species (Soroye et al., 2018). In fact, several studies have been published in recent years using citizen science databases as main data source (e.g. Dennis et al., 2017; Taylor et al., 2019; Barve et al., 2020; White et al., 2023). In fact, the Global Biodiversity Information Facility (GBIF) has been integrating select citizen-science data sources as an additional information source since 2012 (GBIF, 2023).

We used *Lupinus angustifolius* (blue lupine or narrow-leaved lupine) as a study species to explore the potential role of climate change on the evolution of Mediterranean plant phenology. It is a widespread species, easily recognizable and with a large amount of data stored in herbaria, but also present in other data sources, including citizen science databases. The objectives of our study are (1) to evaluate the changes in flowering phenology of *L. angustifolius* populations in the Iberian Peninsula using different data sources, (2) to study potential shifts in vernalization and thermal time of *L. angustifolius* populations using the same data sources, and (3) to assess the differences and level of effectiveness of these historical and modern data sources.

## MATERIAL AND METHODS

### Study species

*Lupinus angustifolius* L. (Fabaceae) is a self-pollinated annual herbaceous plant distributed mainly throughout the Mediterranean Basin, preferring well-drained, sandy, slightly acidic to neutral soils, with a pH in the range of 5 to 7.8. This preference primarily limits its distribution to the western part of the Iberian Peninsula, although it can also be found in some areas with acidic substrates in the east and northeast of the region. The western half of the Iberian Peninsula is considered the center of genetic diversity of the species, where it is most widely distributed (Mousavi-Derazmahalleh et al., 2018).

*L. angustifolius* has palmate leaves with narrow leaflets, and its racemose inflorescence contains bluish-purple flowers. Under natural conditions, *L. angustifolius* flowers tend exclusively to self-pollinate, as this occurs in initial stages of development when the flower is still closed (Kazimierska & Kazimierski, 2002; Wolko et al., 2011). The flowering and fructification season takes place between March and August (Castroviejo & Pascual, 1999).

### Flowering data acquisition

The acquired data consisted of two source types: herbarium specimens and field-taken photographs. A first group of herbarium specimens comprised digitized specimens available in the Global Biodiversity Information Facility (GBIF) (https://www.gbif.org/). A search was made filtering by species, records of the Iberian Peninsula with available image, from 1800 to 2023, and exclusively herbarium entries (https://doi.org/10.15468/dl.4k2qc6). To complement this collection, herbaria holding specimens from populations in the Iberian Peninsula were also contacted, because the digitization of herbarium specimens and uploading to GBIF has not been completed for all Ibero-Macaronesian herbaria. Consequently, we were able to review the material directly or obtain images of the vouchers that had not yet been uploaded to GBIF. To highlight the information not yet available in GBIF, we noted “GBIF” for the first dataset and “Herbarium” for the second. For each record, the herbarium identification code, source herbarium, URL address of the revised voucher, locality, and province to which the municipality belongs, latitude and longitude coordinates (decimal data, in WGS84), and date of collection of the sheet were noted.

Field-taken photographs were sourced from the citizen science platform iNaturalist (https://www.inaturalist.org/) and GeoCAT, a geospatial conservation assessment tool (https://geocat.iucnredlist.org/editor, Bachman et al., 2011) that can incorporate data from different databases and social networks. In this case, we again filtered for records from the Iberian Peninsula, from 1950 to 2023, with precise geographic coordinates and assigned research grade. Subsequently, records that corresponded with some stages of flowering were selected.

### Estimation of flowering peak

*L. angustifolius* has an apical inflorescence of indeterminate growth with acropetal maturation, i.e. the basal flowers mature before the apical ones. Thus, in the same inflorescence we can find fruits forming in the basal part and apical flower buds. This species has a typical spring flowering pattern in the Iberian Peninsula: the first flowers appear at the beginning of March, or even at the end of February, and the last ones in July. Fruiting usually ranges from April to August (Castroviejo & Pascual, 1999).

To estimate the time of flowering, the flowering stage of each individual plant was noted. A four-stage categorization was applied: IF-Initiation of flowering (only two or three flowers can be seen on the main branch); PF-Peak flowering (several opened flowers can be seen on the main branch and some on the lateral branches, if any); FT-Tardy flowering: some withered flowers or even some small fruits and opened flowers can be seen towards the apex of the inflorescence; FD-Fructification in development, several fruits of good size and an occasional flowers are observed. According to this classification, we estimated the days to peak flowering (PF) from January 1^st^, applying the following correction for specimens that were found in the other stages: IF (+5 days); PF (+0 days); FT (-5 days); FD (-15 days). These estimates were applied according to the known flowering development time of the species, based on our findings and previous work (Sacristan-Bajo et al. 2023 a, b; Poyatos et al. 2023).

### Estimation of vernalization and thermal time

*L. angustifolius* flower initiation is influenced by thermal time, photoperiod, and vernalization (Edwards et al., 2011). However, advancements in flowering onset observed in our study area suggest photoperiod is not a limiting factor (Roeber et al., 2022). We therefore investigated whether adjustments occurred in the vernalization and temperature accumulation requirements.

To achieve this, we obtained high-resolution (1 km) daily climate data for our study area. The data, spanning 1950 to 2022, was acquired from the European climatic database hosted at the University of Natural Resources and Life Sciences of Vienna (Moreno & Hasenauer, 2016) and accessed through the *easyclimate* package in R (Cruz-Alonso et al., 2023). This database provides daily minimum (T_min_) and maximum (T_max_) temperatures, which we utilized to calculate thermal time and estimate the number of vernalization days for each study year and site. These parameters were estimated from January 1st for each year. Thermal time (Jones 1992) is the sum of daily mean temperatures (T_mean_) calculated as:

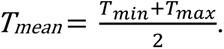

Vernalization days were defined as the number of days with T_mean_ between 1°C and 14°C. These parameters were estimated for each record from January 1st until the date of flowering peak. This approach allowed us to assess how local climatic conditions may be affecting vernalization time and thermal accumulation, crucial for understanding the flowering of *L. angustifolius* in our study area (Edwards et al., 2011).

### Statistical analysis

We assessed flowering time (flowering peak) response in *L. angustifolius* using a generalized linear model (GLM) with a Poisson error distribution and a logarithmic link function to account for potential variations in latitude, elevation, and collection year. By incorporating latitude and elevation as covariates, we aimed to address potential environmental variations across the sampled latitudinal gradient and within the diverse elevations of the Iberian Peninsula. The model also included interactions between year and both latitude and elevation.

Vernalization and thermal time were also explored in separate models, with a Poisson GLM for vernalization and a linear model (LM) for thermal time. Both models included year of collection and the same covariates (year of collection, latitude, and elevation) to assess potential shifts in cold and heat requirements for flowering.

For the GLM and fitted LMs, we assessed goodness-of-fit using R-squared for the linear models and pseudo-R-squared for the GLM, calculated by comparing the residual deviance to the null deviance. Model residuals were checked visually for normality and homogeneity of variance using diagnostic plots (Chatterjee & Hadi 2015). Cook’s distance was employed to identify influential observations within the fitted models. Graphical examination of Cook’s distance values (Ci) was performed to detect outliers, with particularly high Ci values flagged for further scrutiny (Chatterjee & Hadi 2015). Observations with a Cook’s distance exceeding the threshold of 0.5 were subsequently removed to mitigate their impact on model fit. As a result of this screening, two data points were excluded from the flowering time model, and one data point each was removed from the models adjusted for vernalization and thermal time. All models were fitted using the “stats” package in R version 4.2.2 (R Core Team, 2022).

## RESULTS

Total observations were 682, and herbarium data source provided the greatest amount of data, followed by iNaturalist, GBIF, and GeoCAT, with 305, 195, 152, and 30 observations respectively (Figure S1, Table S1). A total of 18 herbaria contributed to the final dataset (Table S1). Data points prior to 1950 were poorly represented, comprising approximately 10% of the complete dataset (Figure S1). Consequently, they were excluded from further analyses. Before 2009 there were only herbarium data available, with or without digitized image in GBIF. From 2009 onwards, records from citizen databases were also obtained. Since then, herbarium records for this species were scarce (Figure S1). It is also noteworthy that only a small percentage of the herbarium sheets have been digitized, Particularly, less than 20% of the records between the 1970s and 2000s, have been scanned and uploaded to GBIF. Regarding the geographical distribution, the data covered the entire distribution area of the species in the Iberian Peninsula (Figure 1).

**Figure 1.**
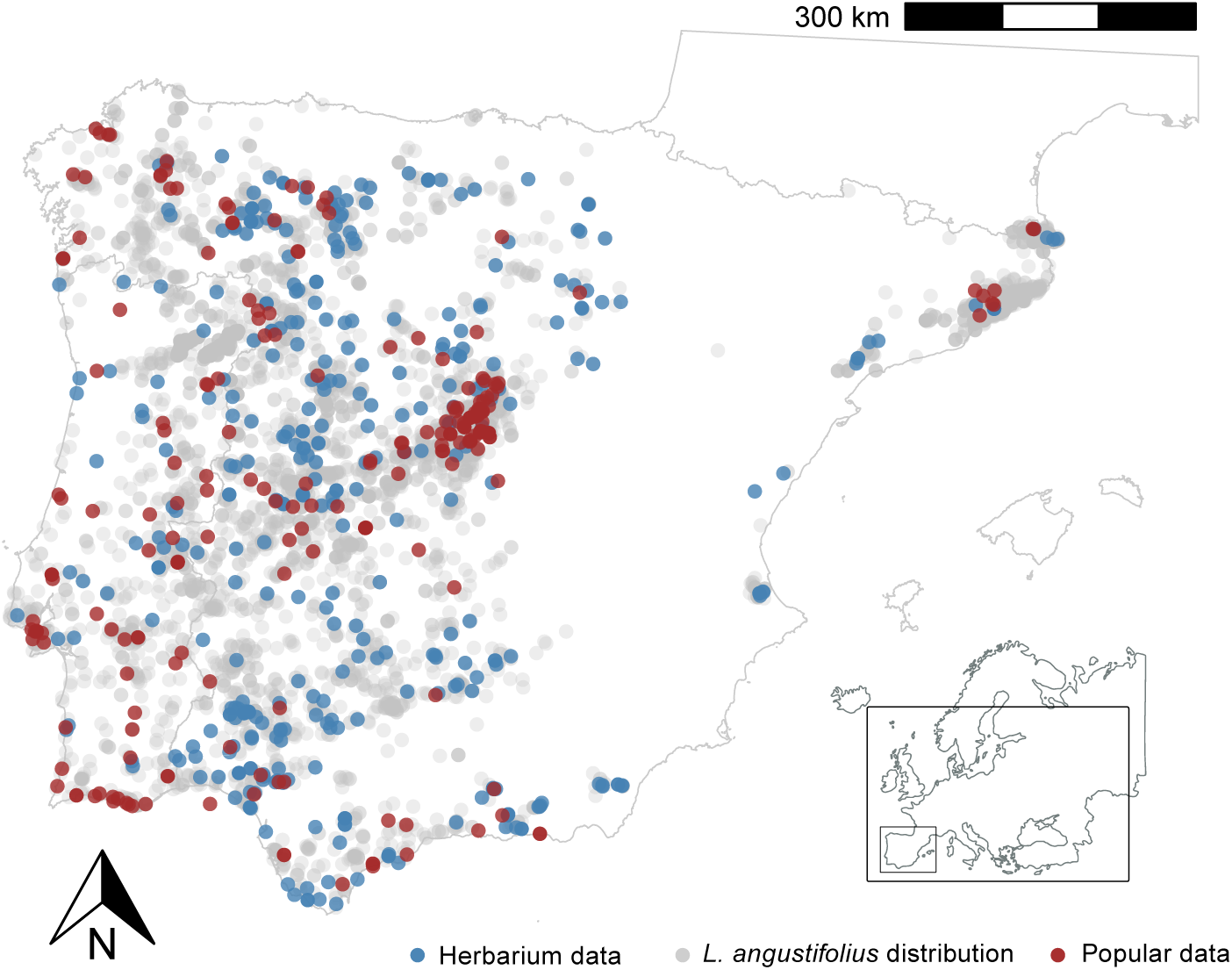
Distribution of *L. angustifolius* across the Iberian Peninsula (in grey), alongside the distribution of flowering observations over the period 1950-2023. Observations from citizen science data sources are highlighted in red, while herbarium records are highlighted in blue.

The GLM fitted for flowering peak revealed that year, latitude, and elevation of collection significantly contributed to explaining the flowering phenology of *L. angustifolius* (all p-values < 0.05, Table 1) with a pseudo-R^2^ of 0.23. A progressive advancement in the flowering of *L. angustifolius* over the study period was observed (Figure 2A). Regarding latitude, plants located farther north flowered later, regardless of the sampled year (Table 1, Figure 2B). A similar pattern was observed with elevation: plants located at higher elevations also flowered later (Table 1, Figure 2C). There were no significant interactions of year with latitude and elevation (Table 1), indicating a similar flowering advance over the years across different latitudes and elevations.

**Figure 2.**
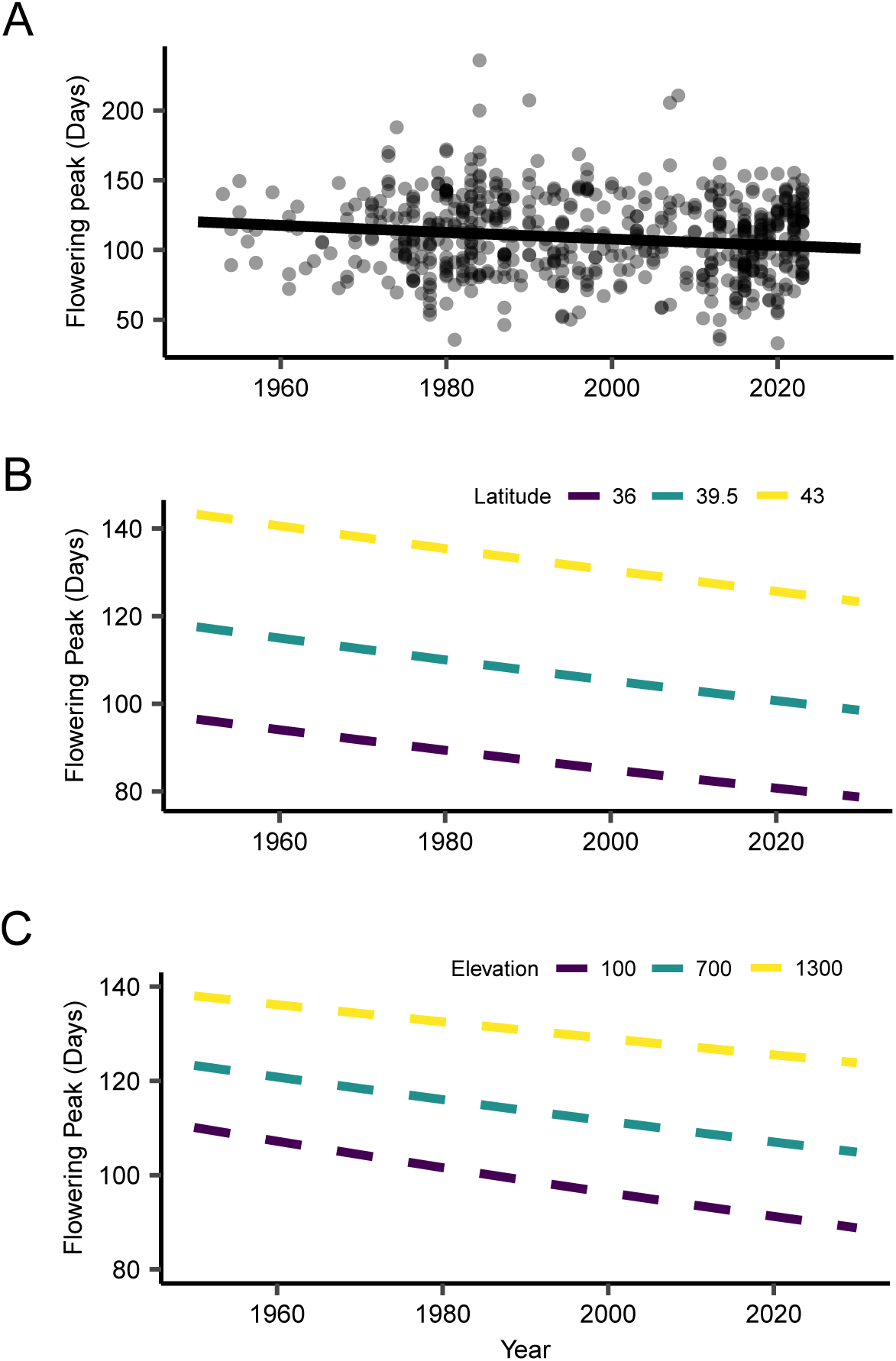
Relationship between sampling year, latitude, elevation, and flowering peak of *L. angustifolius*. (A) Scatter plot showing the relationship between flowering peak and year for observations from 1950 onwards. The points on the graph are slightly jittered horizontally for better visualization. The original data can be found in table S1. The line represents the regression between year and flowering peak. (B, C) Predicted effects of year and latitude or year and elevation on *L. angustifolius* flowering peak from GLM. Dashed lines represent the regression lines for each level of latitude or elevation. Colors represent different latitude or elevation levels.

**Table 1.** Parameter estimates, standard errors, and p-values for the GLM adjusted for the flowering peak of *L. angustifolius*. Continuous predictors are mean-centered and scaled by 1 s.d. The outcome variable remains in its original units. The number of observations was 607, and the model used a Poisson family with a log link function.

| Predictor | Estimate | S.E. | Z val. | *p* |
| --- | --- | --- | --- | --- |
| Intercept | 4.685 | 0.004 | 1159.479 | <0.0001 |
| Year | -0.040 | 0.004 | -9.991 | <0.0001 |
| Latitude | 0.117 | 0.005 | 25.949 | <0.0001 |
| Elevation | 0.085 | 0.004 | 19.712 | <0.0001 |
| Year x Latitude | 0.003 | 0.005 | 0.730 | 0.466 |
| Year x Elevation | 0.007 | 0.004 | 1.713 | 0.087 |

Similarly, the GLM for vernalization (pseudo-R² = 0.40) revealed a progressive advancement over years, indicating that plants are experiencing fewer days of vernalization to flower (Table 2, Figure 3A). Plants located farther north and at higher elevations experienced higher vernalization (Table 2, Figures 3B & 3C). The interaction of year with latitude and elevation was significant (Table 2), indicating a greater decrease in the number of vernalization days at lower latitudes and elevations (Table2, Figure 3B, 3C).

**Figure 3.**
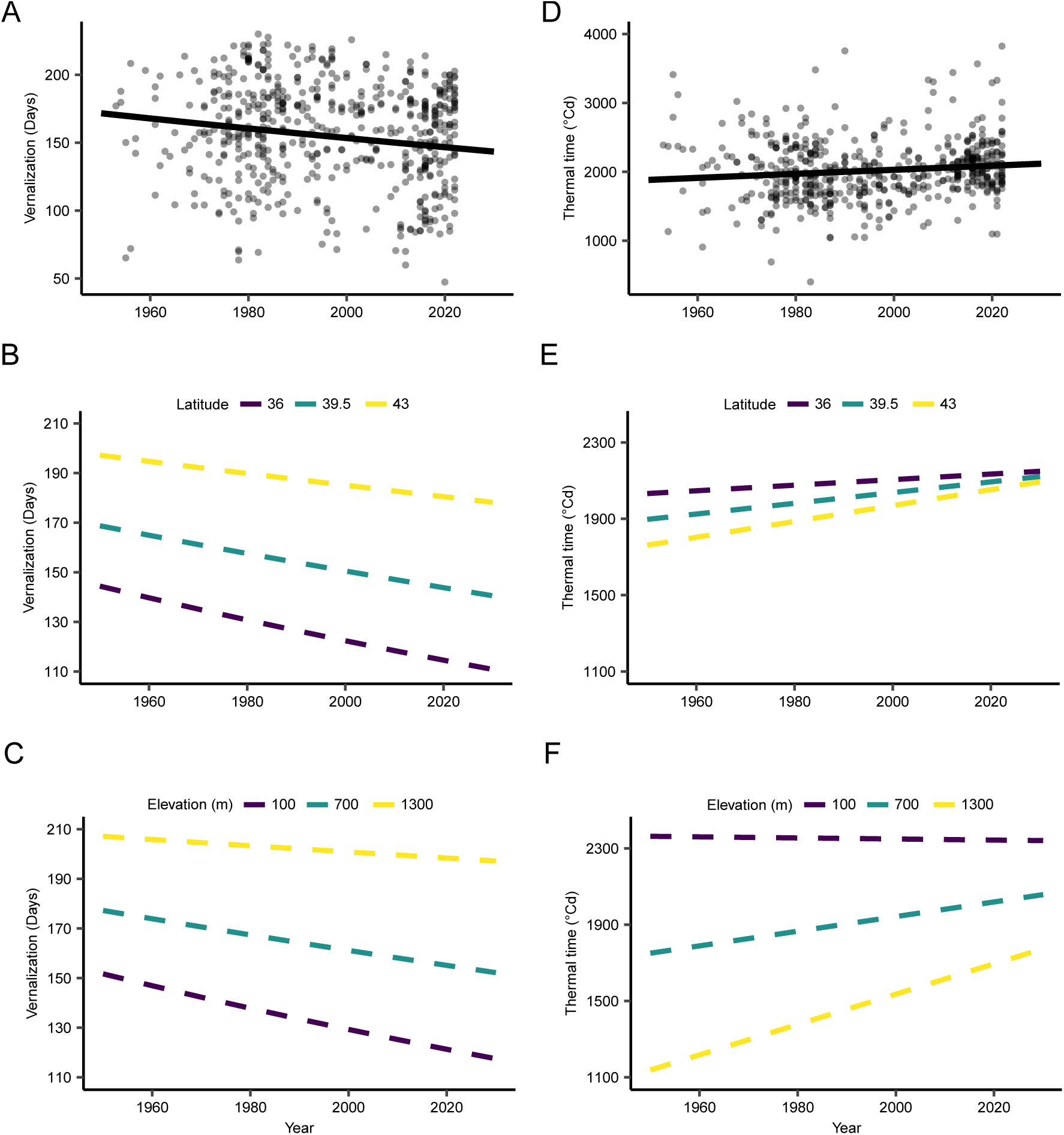
Changes in Vernalization and Thermal Time experienced by *L. angustifolius* populations over time. (A) Scatter plot showing the trend in vernalization days over the years. (B) Modeled trends in vernalization days at different latitudes (36°, 39.5°, and 43°). (C) Modeled trends in vernalization days at different elevations (100 m, 700 m, and 1300 m). (D) Scatter plot showing the trend in thermal time (°C) over time. (E) Modeled trends in thermal time (°C) at different latitudes (36°, 39.5°, and 43°). (F) Modeled trends in thermal time (°C) at different elevations (100 m, 700 m, and 1300 m). Gray dots represent individual observations, and solid lines represent GLMs predictions. Colors represent different latitude or elevation levels.

**Table 2.** Parameter estimates, standard errors and p-values for the model adjusted for the vernalization and thermal time experienced by *L. angustifolius*. Continuous predictors are mean-centered and scaled by 1 s.d. The outcome variable remains in its original units. The number of observations was 607.

| Predictor | Estimate | S.E. | Z val. | *p* |
| --- | --- | --- | --- | --- |
| Vernalization |  |  |  |  |
| Intercept | 5.0413 | 0.0035 | 1455.8858 | <0.0001 |
| Year | -0.0406 | 0.0034 | -11.8341 | <0.0001 |
| Latitude | 0.1099 | 0.0039 | 28.4455 | <0.0001 |
| Elevation | 0.1309 | 0.0037 | 35.3384 | <0.0001 |
| Year x Latitude | 0.0102 | 0.0041 | 2.4625 | 0.0138 |
| Year x Elevation | 0.0093 | 0.0036 | 2.5572 | 0.0106 |
| Thermal time |  |  |  |  |
| Intercept | 2020.2602 | 13.5179 | 149.4503 | <0.0001 |
| Year | 53.1212 | 13.6017 | 3.9055 | 0.0001 |
| Latitude | -39.2927 | 15.2494 | -2.5767 | 0.0102 |
| Elevation | -246.11 | 14.9541 | -16.4577 | <0.0001 |
| Year x Latitude | 13.1829 | 16.1156 | 0.818 | 0.4137 |
| Year x Elevation | 43.4331 | 14.8302 | 2.9287 | 0.0035 |

Thermal time was positively associated with year, indicating an increase in thermal time over time (Table 2, Figure 3D). Latitude and elevation both exhibited a significant negative effect (Table 2, Figure 3E, 3F), implying a decrease in thermal time with increasing latitude and higher elevations. The interaction between year and elevation was significant (Table 2), suggesting a more pronounced increase in thermal time over time at lower elevations. Conversely, the interaction between year and latitude was not significant (Table 2). The model explained 43% of the variation in thermal time (R^2^ = 0.43).

## DISCUSSION

Plant adaptation responses to global change are complex and extremely dynamic, involving multiple traits, genotype-phenotype-environment interactions, and standing population genetic diversity, among others (Exposito-Alonso et al., 2022). Our study reveals that the flowering of *L. angustifolius* has advanced over time, for the last 75 years. This advancement has not been uniform, as populations at lower latitudes and elevations have shown more pronounced advancements in flowering. Additionally, an increase in thermal time has been observed, suggesting a genetic adaptation to flower later, compensating for the earlier flowering induced by rising temperatures. On the other hand, vernalization time has decreased, likely reflecting milder winters due to climate change. This decrease has been more pronounced in populations at lower latitudes and elevations, which could imply stronger natural selection against genotypes requiring vernalization in these areas. These observations are possible thanks to the existence of historical databases, which are useful to studying and evaluating the success of changing strategies and shifting rates (Anderson & Song, 2020).

### Flowering triggered influenced by vernalization and thermal time

Our results may assist in elucidating the underlying mechanisms of the observed earlier flowering in *L. angustifolius*, potentially in response to climate change, as evidenced in other species (Willens et al., 2022). This is partly due to increased thermal time or degree days, where rising temperatures allow plants to reach the thermal thresholds for flowering more quickly (Tang et al., 2016; Piao et al., 2019). This shift has significant evolutionary implications, as plants able to adjust their life cycles are more likely to persist and reproduce. *L. angustifolius* flowering time is heritable and subject to selection (Sacristán-Bajo et al., 2023a, b; Poyatos et al., 2023), making it likely to evolve in response to strong selective pressure, such as climate change (Franks et al., 2007; Franks & Weis 2008; Franks, 2011). Moreover, previous research showed that plasticity can generate adaptive phenology variation in this species (Matesanz et al., 2020), and both genetic differentiation and plasticity are crucial for species adaptation, but the relative importance of the two processes is contingent upon the speed and nature of environmental change, the time and generations involved, and the genetic variability available (Franks & Hoffmann, 2012).

In that sense, our analysis showed uneven phenological shifts across the Iberian Peninsula, with populations at lower latitudes and elevations experiencing different shifts in vernalization and thermal time requirements for flowering. Studies on latitude’s impact on phenology show mixed results, with some suggesting northern plants are less sensitive to temperature changes (Ge et al., 2015; Shen et al., 2015), while others suggest the opposite (Pudas et al., 2008; Prevéy et al., 2017). Degree-day accumulation rates are faster at lower latitudes, accelerating plant development cycles (Visser & Lindner, 2018; Freeman et al., 2021). Similarly, research on elevation effect showed contrasting findings: some indicate greater sensitivity at high elevations (Ziello et al., 2009; Kopp et al., 2020), while others argue the contrary (Defila & Clot, 2005; Cuffar et al., 2012; Willems et al., 2022). This discrepancy likely arises from varying spatial scales and the complex, non-linear relationship between elevation, latitude, and phenology, influenced by other environmental factors (Willems et al., 2022). In our study, elevation and latitude patterns imply well-conserved inverse temperature gradients, which would explain the linear and positive relationship between flowering phenology and both geographical variables. Flowering is earlier at all latitudes and elevations. Populations at colder locations flower later, but there is no interaction with years.

Evidence suggests that thermal time for flowering has increased over time, suggesting a genetic adaptation to flower later, compensating for earlier flowering driven by rising temperatures. This trend is more evident in populations at higher latitudes and elevations, where delayed flowering appears to reflect stronger genetic adaptations to local climatic conditions. At the same time, vernalization requirements have decreased globally, allowing plants to flower earlier by accumulating fewer vernalization days and responding to higher temperatures (Cook et al., 2012; Duncan et al., 2015). In *L. angustifolius*, this reduction in vernalization needs is more pronounced at lower latitudes and elevations, suggesting that these requirements are not fixed but vary among populations based on environmental pressures. Cold-adapted populations, by contrast, show reduced responses to vernalization (Preston & Sandve, 2013), possibly due to decreased phenotypic plasticity as evolutionary protection against early frosts or inactivated physiological mechanisms (Preston & Sandve, 2013). Nevertheless, photoperiod may also be playing a crucial role here, with stronger responses observed along latitudinal gradients than elevational ones. Even when vernalization and temperature thresholds are met, insufficient light can prevent flowering (Mouradov et al., 2002). This suggests that photoperiod sensitivity could serve as an additional mechanism to avoid early flowering and late frosts (Inouye, 2000). The accumulation of degree days further indicates that some populations require cold periods even after reaching minimum temperature thresholds, emphasizing the complexity of flowering cues. These adaptations, particularly the reduction in vernalization needs, are likely driven by climate change and milder winters. Genotypes requiring vernalization face increasing disadvantage, especially in regions where winter warming is more pronounced, such as lower latitudes and elevations (Hansen et al., 2012; Mountain Research Initiative EDW Working Group, 2015). Therefore, at higher latitudes and elevations, the slower advance in flowering could reflect adaptations to avoid late frosts, leading to greater reliance on photoperiod (Renner & Zohner, 2018), although further research is needed to confirm these hypotheses.

### Data quality and database biases

Our results show that both herbarium records and field-taken photo database significantly contribute to plant phenology studies, particularly to flowering phenology, as flowers are more often photographed and hold greater taxonomic relevance. Over time, differences in record types have become clear. For example, the number of herbarium records has drastically declined, a trend noted in prior research (Prather et al., 2004a, b). Despite this decline, herbarium records remain useful for understanding reproductive phenology patterns (Cleland et al., 2007; Jeong et al., 2011; Fu et al., 2014), though newer tools like iNaturalist are gradually supplementing this role (Meineke et al., 2014; Zhou et al., 2022). Symptomatic of this is the number of *L. angustifolius* records observed on iNaturalist during the last decade (Figure S1), which comprises half of the data from our analysis in a time fraction equivalent to one-fifth of the total period analyzed.

Regional and temporal discrepancies in collection efforts mean that data are sometimes insufficient for robust conclusions (Magurran & Hendriks, 2003; Benayas et al., 2009). Increased collection efforts and better integration into databases like GBIF, which centralizes data from multiple sources such as iNaturalist and GeoCat, could enhance our understanding of plant dynamics. However, we observed a decrease in herbarium records in GBIF, likely due to either outdated data uploads or reduced field collection efforts. It’s crucial to continue collecting herbarium specimens, as they serve as important spatio-temporal records of individual plants (Benayas et al., 2009). Although modern tools like iNaturalist offer easy access to large datasets, they come with challenges, including taxonomic misidentifications (Contreras-Diaz et al., 2023). These errors, often due to incomplete photos or automatic identification software, underscore the importance of physical herbarium records for precise taxonomic work. For example, our study of *L*. *angustifolius* involved manually reviewing all available photos to eliminate confusion with similar species like *Lupinus consentinii* and *Lupinus micranthus.* This differentiation was based on key characteristics such as the width of the leaflets and the insertion of the flowers along the scape (Castroviejo & Pascual, 1999), ensuring accurate species identification amidst potential errors in automated systems.

In sum, while each data source has strengths and weaknesses, their complementarity is undeniable. It is up to the scientific community to carefully filter and validate these datasets (Lopez-Guillen et al., 2024) to ensure that conclusions are drawn from reliable data sources.

## Acknowledgements

This publication is part of the R&D&I project PID2021-127841OA-I00 funded by MICIU/AEI /10.13039/501100011033/ and by ERDF A Way for Europe, coordinated by CL and AG; and by the grant FJC2020-044244-I awarded to FFJL, funded by MCIN/AEI /10.13039/501100011033 and NextGeneration EU/PRTR. The authors would like to express their gratitude to the herbaria ARAN, BC, GDA, LEB, SALA, SEV, and VAL for providing their specimens for consultation, as they were not available online through GBIF.

