## Supplementary material for "Evidence of flowering time advance in blue lupin (*Lupinus angustifolius*) in the last decades revealed by herbarium data and citizen science databases": Supplemetary Files

**SUPPLEMENTARY INFORMATION**

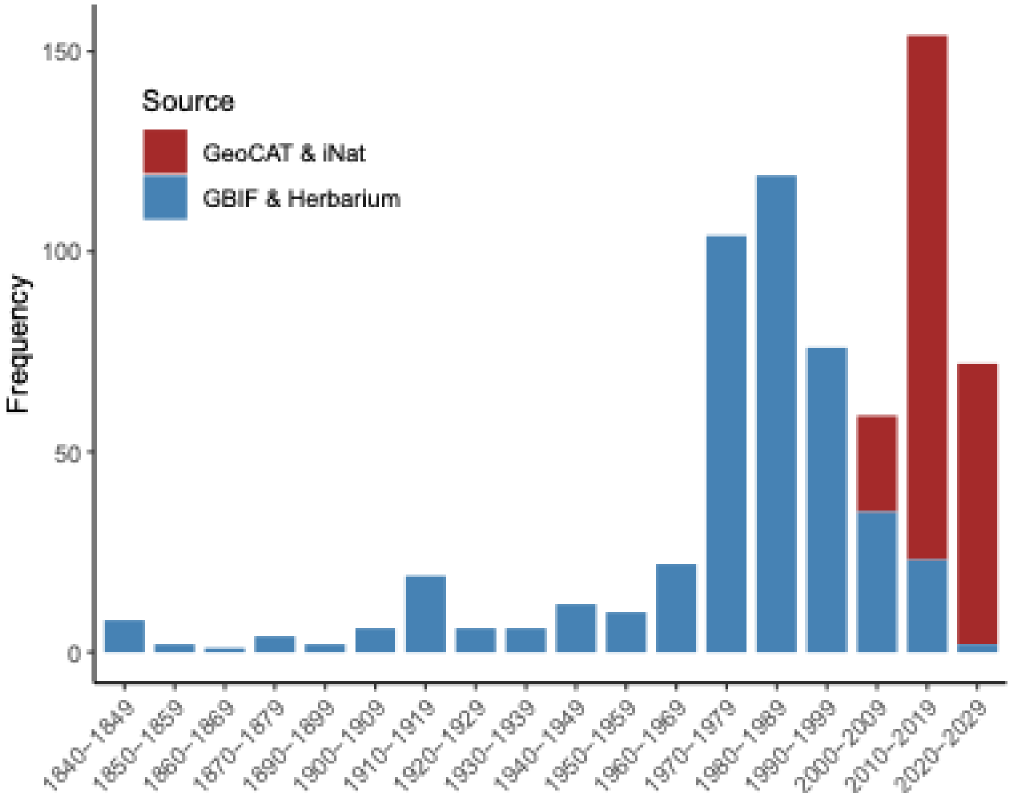


**Figure S1.** Chronological distribution of records by source over time of flowering phenology data for *Lupinus angustifolius* in the Iberian Peninsula. The data are grouped into two categories: "GeoCAT & iNat," representing citizen science records obtained from platforms such as iNaturalist and GeoCAT, and "GBIF & Herbarium," which includes digitized herbarium specimens and data from GBIF. The frequency of records per decade shows a significant increase in observations in the last decade, especially in the citizen science category, highlighting the importance of these platforms for phenological monitoring in recent years.

**Table S1.** Summary of herbarium records or citizen science data grouped by institution for *Lupinus angustifolius* in the Iberian Peninsula. Herbarium codes follow the standard from the *Index Herbariorum*, and citizen science records from iNaturalist and GeoCAT are listed separately. This table highlights the contribution of both traditional herbarium data and modern citizen science platforms to the phenological monitoring of the species.

| Herbarium | Source | Number of Records | Earliest Year | Latest Year | Total Online Records | Percentage Online Records |
|---|---|---|---|---|---|---|
| ARAN- Aranzadi Zientzia Elkartea (Spain) | Herbarium data | 7 | 1984 | 2004 | 0 | 0.0 |
| LEB - Universidad de León (Spain) | Herbarium data | 48 | 1969 | 2016 | 0 | 0.0 |
| SALA - Universidad Salamanca (Spain) | Herbarium data | 54 | 1964 | 2020 | 0 | 0.0 |
| SEV - Universidad de Sevilla (Spain) | Herbarium data | 85 | 1907 | 2019 | 0 | 0.0 |
| BC - Institut Botànic de Barcelona (Spain) | Herbarium data | 18 | 1872 | 2022 | 0 | 0.0 |
| COI - Herbarium of the University of Coimbra | Herbarium data | 21 | 1851 | 2015 | 21 | 100.0 |
| VAL - Universitat de València (Spain) | Herbarium data | 20 | 1968 | 2022 | 0 | 0.0 |
| LSU - Louisiana State University (USA) | Herbarium data | 1 | 1977 | 1977 | 1 | 100.0 |
| BR - Meise Botanic Garden (Belgium) | Herbarium data | 10 | 1849 | 2005 | 10 | 100.0 |
| GDA - Universidad de Granada (Spain) | Herbarium data | 29 | 1967 | 2016 | 0 | 0.0 |
| P - Muséum National d'Histoire Naturelle (France) | Herbarium data | 7 | 1849 | 2016 | 7 | 100.0 |
| US - Smithsonian Institution (USA) | Herbarium data | 1 | 1848 | 1848 | 1 | 100.0 |
| WUP - University of Vienna (Austria) | Herbarium data | 2 | 2013 | 2016 | 2 | 100.0 |
| L - Naturalis Biodiversity Center (The Netherlands) | Herbarium data | 3 | 1980 | 1985 | 3 | 100.0 |
| MA - Real Jardín Botánico (Spain) | Herbarium data | 144 | 1856 | 2016 | 144 | 100.0 |
| O - University of Oslo (Norway) | Herbarium data | 1 | 1957 | 1957 | 1 | 100.0 |
| SANT - Universidad de Santiago de Compostela | Herbarium data | 1 | 2005 | 2005 | 1 | 100.0 |
| GeoCAT | Citizen science | 30 | 2006 | 2023 | 30 | 100.0 |
| iNaturalist | Citizen science | 195 | 2002 | 2023 | 195 | 100.0 |
